# An algal nutrient-replete, optimized medium for fast growth and high triacylglycerol accumulation

**DOI:** 10.1101/2024.11.07.622501

**Authors:** Tim L Jeffers, Ryan McCombs, Stefan Schmollinger, Sabeeha S Merchant, Krishna K Niyogi, Melissa S Roth

## Abstract

Microalgae are promising sources to sustainably meet the global needs for energy and products. Algae grow under different trophic conditions, where nutritional status regulates biosynthetic pathways, energy production, and growth. The green alga *Chromochloris zofingiensis* has strong economic potential because it co-produces biofuel precursors and the high-value antioxidant astaxanthin while accumulating biomass when grown mixotrophically. As an emerging reference alga for photosynthesis, metabolism, and bioproduction, *C. zofingiensis* needs a defined, optimized medium to standardize experiments during fast growth. Because the interplay of glucose consumption (+Glc) and mineral deficiency influences photosynthesis, growth, and the production of lipids and astaxanthin, we designed a replete nutrient medium tailored to the *C. zofingiensis* cellular ionome. We combined inductively coupled plasma mass spectrometry (ICP-MS) and +Glc growth curves to determine a medium that is nutrient replete for at least 5 days of +Glc logarithmic growth. We found that there are high nutritional needs for phosphorus and sulfur during mixotrophy. Iron was the only element measured for which the cellular concentration correlated with exogenous concentration and was iteratively adjusted until the internal ionome was consistent through the logarithmic growth phase. This *<u>C</u>hromochloris*-<u>O</u>ptimized <u>R</u>atio of <u>E</u>lements (CORE) medium supports fast growth and high biomass without causing excess nutrient toxicity. This defined, nutrient-replete standard is important for future *C. zofingiensis* investigations and can be adapted for other species to support high biomass. The method used to develop CORE medium shows how ionomics informs replicable media design and may be applied in industrial settings to inform cost-effective biofuel production.

**Significance Statement:** Studying how carbon sources and mineral nutrients interplay to regulate algal metabolism can be exploited to discover and control pathways in photosynthesis and biofuel production. Here we design a medium from the cellular ionome of *Chromochloris zofingiensis*, a powerful algal model for photosynthesis, metabolism, and bioproducts, to provide a defined, replete standard for mixotrophic and heterotrophic growth of green algae. These media design principles show how accounting for increased nutritional demands based on carbon substrate can ensure experimental replicability when probing diverse algal metabolisms.

## Introduction

As global demands for energy and products rise, microalgae are gaining interest as components of a sustainable bioeconomy. Several algae species accumulate high quantities of lipids, making them relatively carbon-neutral sources to replace fossil fuels (Unkefer et al., 2017; Arora et al., 2018). While many algae accumulate the biofuel precursors triacylglycerols (TAGs) under stress conditions at the expense of biomass (Wijffels and Barbosa 2010; Ma et al., 2022), some algae can amass high amounts of TAGs and biomass concurrently (Roth et al., 2019a; Jeffers et al., 2024). These algae are often mixotrophic, capable of both photosynthesis and organic carbon consumption for energy and biomass production (Suzuki et al., 2018; Blaby-Haas and Merchant, 2019). Taking advantage of algal mixotrophy can benefit the industrial cultivation of algae to produce lipids. For example, treating large-scale cultures with reduced carbon and nutrients via wastewater can reduce production costs (Ma et al., 2022). Furthermore, microalgae that produce high-value compounds such as pharmaceuticals, nutraceuticals, and food supplements can improve the economic viability of biofuel through co-production of valuable products. However, a growth medium that can remain nutrient replete to maximize biomass and biofuel and bioproduct accumulation while minimizing nutrient deficiency is necessary for industry and research.

The unicellular green alga *Chromochloris zofingiensis* is an emerging reference alga for research on photosynthesis, metabolism, and bioproduction (Roth et al., 2017; Roth et al., 2019a; Zhang et al., 2021; Wood et al., 2022). Not only is *C. zofingiensis* one of the highest algal producers of TAGs (Breuer et al., 2012), but it also co-produces the high-value nutraceutical astaxanthin (Roth et al., 2017; Jeffers and Roth 2021; Zhang et al., 2021). *C. zofingiensis* has a high-quality complete genome that serves as a foundation for ‘omics studies and accelerates bioproduct pathway discovery (Roth et al., 2017). Supplying sugars to *C. zofingiensis* cultures supports accumulation of lipids, astaxanthin and biomass concurrently (Roth et al., 2019a; Jeffers et al., 2024). Lipid accumulation during enhanced growth contrasts to the more widely studied phenomenon where nutrient deficiency induces storage lipids at the expense of biomass (Arora et al., 2018; Ma et al., 2022). *C. zofingiensis* has evolutionarily distinct mechanisms of photosynthetic regulation and lipid accumulation compared to the reference green alga *Chlamydomonas reinhardtii* possibly in part due to its ability to consume a wider array of carbon sources (Sun et al., 2008; Roth et al., 2017; Suzuki et al., 2018; Jeffers et al., 2024).

*C. zofingiensis*, like many algae, consumes exogenous sugars such as glucose (Glc) (Suzuki et al., 2018; Roth et al., 2019a). In contrast, *C. reinhardtii* cannot consume Glc, although it can be grown with acetate (Salomé and Merchant, 2019). +Glc switches off *C. zofingiensis* photosynthesis and induces lipid accumulation via *de novo* fatty acid synthesis by a hexokinase-dependent pathway (Roth et al., 2019a; Roth et al., 2019b; Jeffers et al., 2024). Recently, we discovered that switching off photosynthesis, but not lipid accumulation, is prevented by treating +Glc cultures with a replete iron supplement (Jeffers et al., 2024). The switching off of photosynthesis in the light only occurs when iron is limiting but not with other nutrient depletions (Jeffers et al., 2024). These studies were conducted in Proteose medium (Bristol’s + Proteose peptone + Chu’s micronutrient medium), which is a rich and undefined growth medium with low iron content (Roth et al., 2019a; Roth et al., 2019b; Jeffers et al., 2024). Previous studies with *C. zofingiensis* have also used a modified Bristol’s medium (Sun et al., 2008) and Kuhl medium (Huang et al., 2016; Zhang et al., 2017). Regardless of the medium, adding Glc substantially improves biomass and TAG accumulation. As nutrient deficiency has unique metabolic impacts depending on +Glc vs. -Glc, it is imperative to design a standard *C. zofingiensis* medium to achieve a replete mineral composition as an experimental control for optimal +Glc growth and metabolism.

While nutritional status reshapes algal metabolism, it can be difficult to identify due to “invisible” signatures of nutrient deficiency. For example, in *C. reinhardtii* high-affinity iron transport systems are activated in the iron-deficiency regime (1-3 µM Fe) without any obvious phenotypic changes in chlorophyll content or quantum efficiency of PSII (Merchant et al., 2006; Glaesener et al., 2013). Replete metal nutrition in *C. reinhardtii* was defined through an experimental pipeline where a micronutrient supplement composition was derived from the internal ionome of *C. reinhardtii* cultures measured through ICP-MS (Kropat et al., 2011). The final supplement is three-fold higher than the nutrient content of cellular biomass at stationary phase (Kropat et al., 2011), ensuring cells remained nutritionally replete with only slight excess of nutrients above their maximum biomass needs. At these concentrations, the ionome is consistent through logarithmic growth phase, indicating neither deficiency nor hyperaccumulation from excess impacts cell physiology in this optimized medium (Kropat et al., 2011; Hui et al., 2022).

Here, we employed media design principles based on the cellular ionome to ensure replete concentrations of nutrients were optimized to the internal ratio of elements in *C. zofingiensis* batch-culture experiments. To account for the large increase in biomass in +Glc cultures of *C. zofingiensis*, we enhanced macronutrient concentrations and adjusted for the trade-off between excess nutrient toxicity in photoautotrophy and the high nutrient demands of +Glc cultures. Our final medium, *<u>C</u>hromochloris*-<u>O</u>ptimized <u>R</u>atio of <u>E</u>lements (CORE), was both sufficient for high growth in +Glc yet maintained a consistent internal ionome during photoautotrophic logarithmic growth. This strategy shows how ionome measurements of Glc-consuming algae can inform the experimental design for research replicability and how ionome techniques can be applied in future studies to budget mineral nutrient input with lipid production output in algal bioprospecting systems, facilitating cost-competitive alternatives to fossil fuels.

## Results

### Glucose drives fast growth and high biomass accumulation, but demands more nutrients

Because +Glc induces high TAG and biomass accumulation in *C. zofingiensis* (Roth et al., 2019a; Roth et al., 2019b; Jeffers et al., 2024), we designed a defined, minimal medium that compensated for the increased nutrient budget of Glc-fed batch cultures to keep cells nutrient replete regardless of trophic state. Our previous studies used “Proteose” medium (Roth et al., 2017; Roth et al., 2019a; Roth et al., 2019b; Jeffers et al., 2024), which is composed of Bristol’s medium for macronutrients (Bold, 1949), Chu’s micronutrient supplement for metal nutrients (Chu et al., 1975), and a Proteose Peptone supplement (See Table S1 for this and all media compositions). Proteose provides an additional, undefined source of micronutrients and N-rich organic compounds. However, our previous results showed that a +Fe supplement (≥10 µM Fe) rescues photosynthesis and further increases biomass of +Glc cultures (Jeffers et al., 2024), indicating that the undefined amount of iron in Proteose (estimated between 1-2 µM, M. Meagher pers. comms. 2024) was at a limiting concentration. To make a defined medium, we removed the Proteose Peptone supplement and increased the micronutrient concentrations from Chu’s supplement (Table S1), using similar stock sources and concentrations as optimized for *C. reinhardtii* (Kropat et al., 2011).

Next we increased macronutrients based on the physiological response in +Glc and preliminary spent medium measurements. For example, in replete Fe medium, increasing NO_3_^−^ caused cultures to remain green after +Glc addition (Figure S1), while similar cultures with replete Fe and NO_3_^−^ concentrations near Bristol’s medium levels (3 mM) turned orange or brown with +Glc (Roth et al., 2019b; Jeffers et al., 2024). Compared to Bristol’s, NO_3_^−^ was increased by 20 mM. In addition, preliminary ICP-OES measurements on spent +Glc medium suggested that +Glc increases the sulfur and phosphorous consumption of cultures. Therefore, S and P concentrations were also increased. These modifications to the growth medium generally improved nutrient sufficiency to match the demands of +Glc, and we labelled this medium ADJ (for “adjusted”) (Table S1). With these nutrient increases, ADJ medium became an appropriate starting point to fine tune *C. zofingiensis* medium components according to its general internal ionome needs and biomass levels in response to +Glc.

The media design principles of Kropat et al., 2011 use maximum biomass (i.e., stationary phase) and the cellular ionome to calculate the required mineral nutrient composition for a replete growth medium. To determine a theoretical biomass maximum of Glc resupply experiments, we aimed to see how high *C. zofingiensis* biomass becomes when Glc is not limited in batch cultures. To do so, 20 mM Glc was first added daily to logarithmic growth phase cultures (day 5). Glc concentration in spent medium was measured daily to determine when Glc re-supply needed to be increased to match consumption. Glc treatment immediately increased logarithmic growth rate between days 5-11 (Figure 1A,C) and Glc resupply was 60 mM day^-1^ by day 11 (Figure 1D). Due to continued growth and high Glc consumption, a second experimental phase involved resuspending the cultures in fresh medium with 500 mM Glc every 2-3 days. Cultures continued growth in a second logarithmic growth phase (Figure 1A) that had a lower biomass doubling rate. The experiment continued until day 24, where cellular volume was approximated as ∼22.5% of total liquid culture volume (Figure 1B). Volumetric biomass was approximately 60-fold higher than the stationary biomass of photoautotrophic cultures.

**Figure 1.**
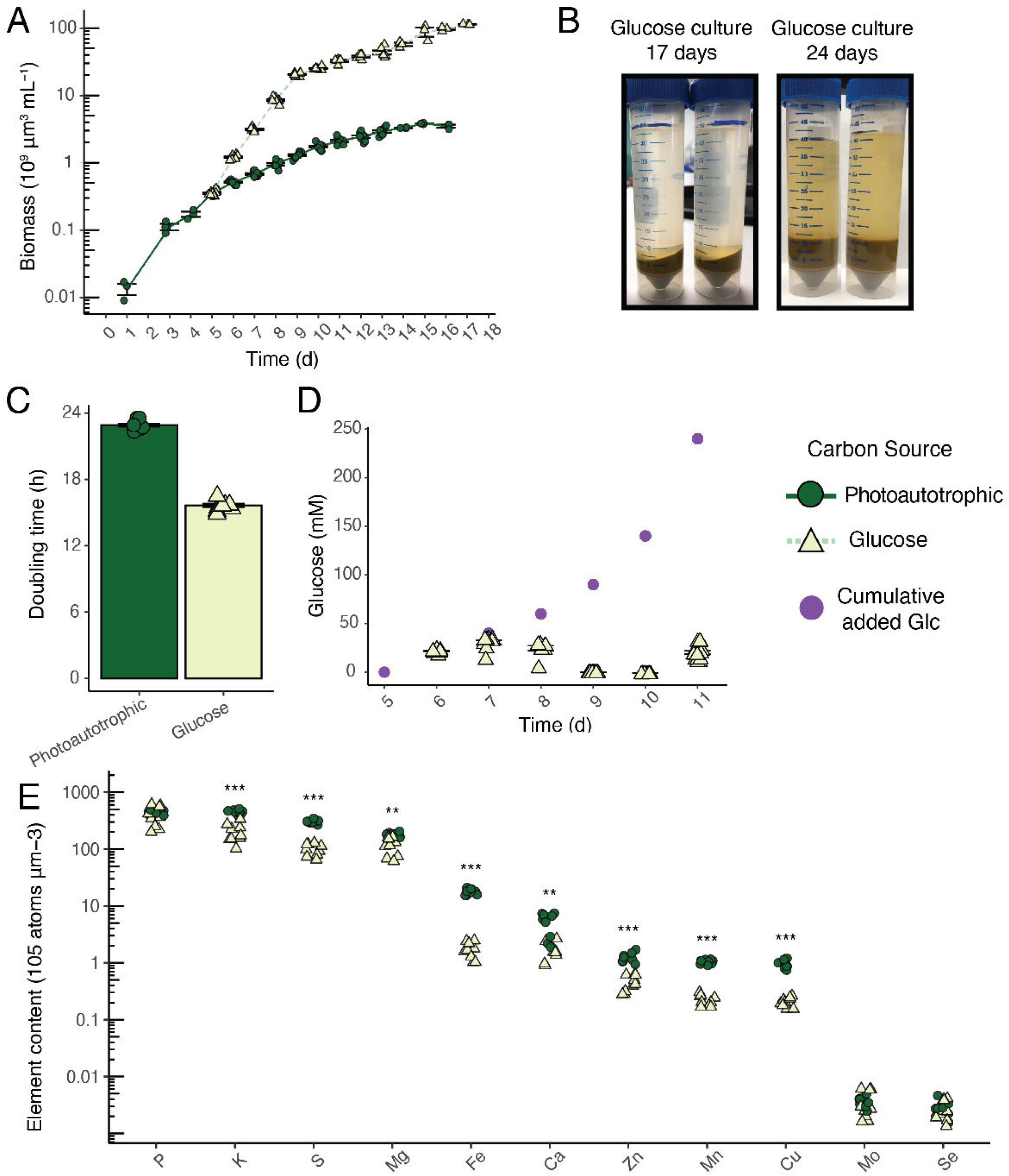
Glucose-fed *C. zofingiensis* has fast, long-term growth accumulating significant biomass. **(A)** Volumetric biomass growth curves of batch cultures of photoautotrophic (pale green, circles, solid lines) and +Glc cultures (dark green, triangles, dotted line) in ADJ medium (Table S1). **(B)** Photographs of centrifuged +Glc cultures after 17 days and 24 days continued maintenance of +Glc and nutrients through resupply. **(C)** Volumetric biomass doubling time (h) of photoautotrophic log phase (calculated between days 1-6) growth and the first +Glc stage of log phase growth (days 5-8). **(D)** Glucose consumption during glucose resupply of batch experiments. The calculated addition of leftover glucose and added glucose is shown in purple. **(E)** Elemental content measured by ICP-MS presented as atoms per volumetric biomass of autotrophic vs. +Glc cells. Asterisks indicate level of significance of student’s *t*-test of the concentration difference between +Glc vs. photoautotrophic cells (*p* < 0.001 ∼ ***, *p* < 0.01 ∼ **, *p* < 0.05 ∼ *). Both the 10^th^ day and 14^th^ day ICP-MS samples were combined for *t*-test analysis and plotting. Line and bar graphs represent means and error bars represent standard error (*n* = 3-11).

The extreme biomass density produced when Glc is not limiting showed a promising industrial possibility of growing high density algal slurries with lower water volume needs. However, for laboratory studies trying to achieve these maximum biomass levels may create unrealistically high nutrient compositions.

### *Identifying the internal ratio elements of* C. zofingiensis *cells*

The internal composition of elements of cells can be multiplied by a target biomass to calculate a replete nutrient budget (Kropat et al., 2011). To measure the ionome, cellular concentrations of P, K, S, Mg, Ca, Fe, Mn, Zn, Cu, Se, and Mo were determined by ICP-MS in -Glc and +Glc cultures on day 10 and 14 of the *C. zofingiensis* growth curve (Figure 1E). Most nutrients displayed significantly lower concentrations per biomass in +Glc vs. -Glc at both timepoints (*p* < 0.05) except for P, Mo, and Se, for which +Glc vs. -Glc concentrations were not significantly different. We suspect the decrease of ionic nutrients per volumetric biomass is due to carbon dense starch and lipid accumulation dominating cellular space in +Glc (Roth et al., 2019a; Jeffers et al., 2024).

In addition, while Ca can compose 0.1 to 5% of vascular plants plant dry weight (Thor, 2019), we observed that Ca levels were lower than Fe levels in *C. zofingiensis*, despite Fe being typically defined as a micronutrient and Ca defined as a macronutrient (Figure 1E). This measurement of low Ca contrasts with the reference green algal *C. reinhardtii*, where Ca is more abundant than Fe in standard conditions where Fe does not hyperaccumulate (Merchant et al., 2020; Schmollinger et al., 2021; Hui et al., 2022). These comparative results indicate *C. zofingiensis* has lower Ca requirements relative to other photosynthetic organisms.

### Optimizing nutrient ratios improves glucose-fed growth

We tested the trade-off between nutrient excess toxicity vs. insufficiency by creating scalar multipliers of the ionome and observing the impacts on algal growth. New media recipes were derived by multiplying the mean elemental concentration per biomass in -Glc by the following growth stages biomasses: <u>p</u>hotoautotrophic stationary stage at day 14 (called P14, the lowest nutrient composition), <u>m</u>ixotrophic (+Glc) at day 10 (M10), and <u>m</u>ixotrophic (+Glc) at day 14 (M14, highest nutrient composition) (Figure 2A, Table S1 for media recipes). For M10, we additionally designed a medium (M10.Ca) that had two-fold higher Ca concentration to ensure that the previously measured low Ca (Figure 2A) was not a result from a nutrient deficiency in ADJ medium. Like the media optimization for *C. reinhardtii*, this composition was multiplied by three for mild nutrient excess and to maintain a similar nutrient environment during logarithmic growth (Kropat et al., 2011). However, unlike for *C reinhardtii*, macronutrient levels also required adjustments to compensate for +Glc biomass increases. Standard ICP-MS protocols use nitric acid for digestion of cell material, preventing the accurate determination of the cellular N content. The NO_3_^−^ concentration was therefore controlled to be at the same ratio to all other elements across tested media. In +Glc-derived M10 and M14 medium, all macronutrients and micronutrients (except Ca) exceeded the composition of revised *C. reinhardtii* TAP medium with micronutrient supplement (Kropat et al., 2011) and the originally used ADJ medium, suggesting that +Glc biomass levels greatly enhance nutrient budget demands (Figure 2A).

**Figure 2.**
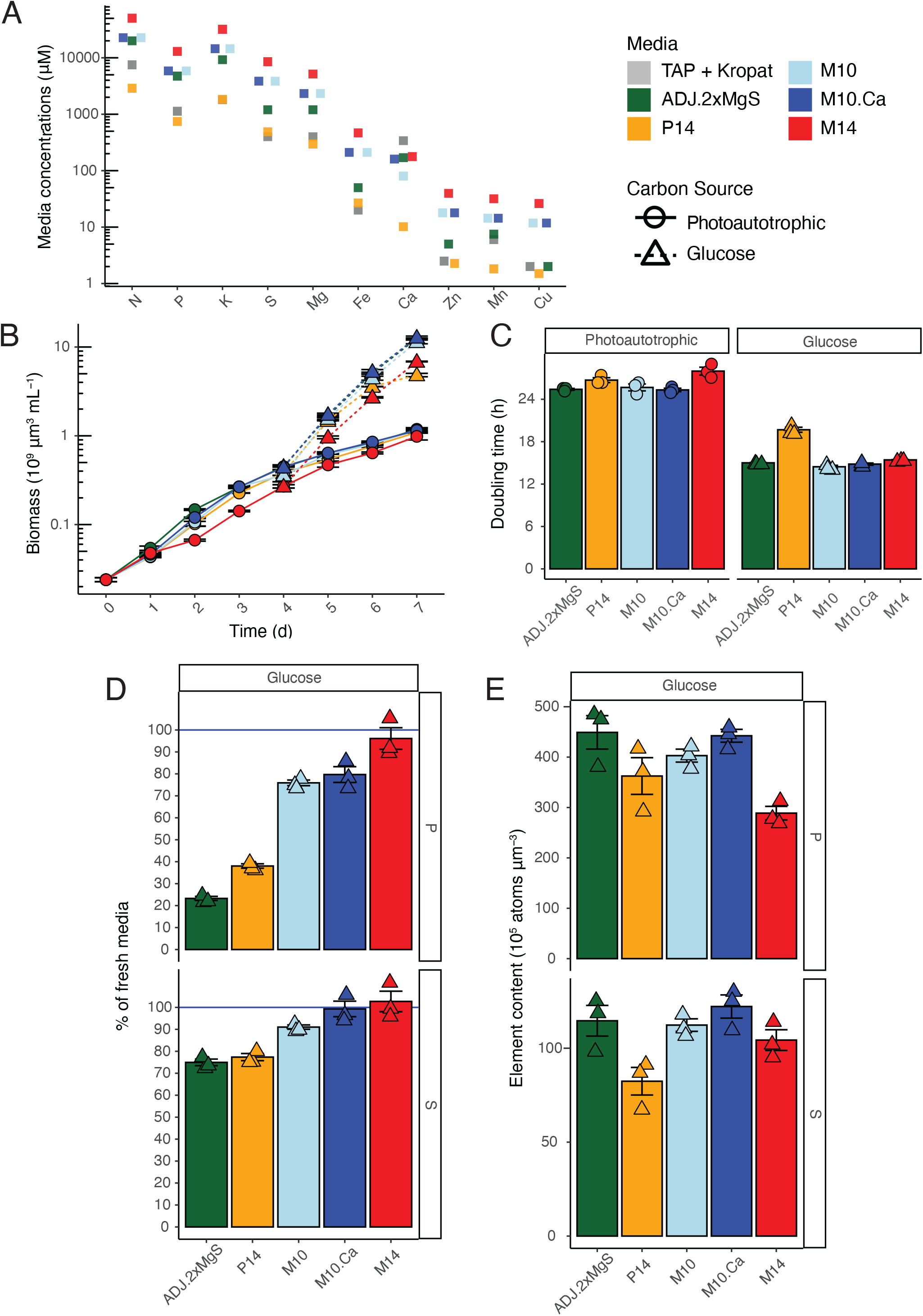
Nutrient concentration trade-offs for photoautotrophic and long-term +Glc growth. **(A)** Elemental nutrient concentrations of media designed to meet nutrient demands of different stages of growth: photoautotrophic (P14, orange) and long-term glucose growth (M10, light blue; M10.Ca, blue; M14, red) (see Table S1 for details). Comparison media for reference are the previously described ADJ medium (green) and micronutrient-optimized TAP medium for *C. reinhardtii* (Kropat et al., 2011, grey). **(B)** Volumetric biomass growth curve of *C. zofingiensis* in media ADJ, P14, M10, M10.Ca, and M14 for photoautotrophic (solid line, circles) and +Glc treatment (dotted line, triangles). Each point is mean values instead of raw due to high overlap in growth rates. **(C)** Volumetric biomass doubling time (h) across media and trophic states. Points are individual growth curves. **(D)** Percentage of sulfur (S) and phosphorus (P) left in spent medium relative to fresh medium in the +Glc condition on day 7 (3 days of +Glc growth). Concentrations are measured by ICP-MS. **(E)** Internal elemental content of S and P presented as atoms per volumetric biomass in day 7 +Glc cells. Line and bar graphs represent means and error bars represent standard error (*n* = 3).

-Glc and +Glc growth curves across media revealed a trade-off between nutrient limitation and excess nutrient toxicity (Figure 2). Photoautotrophic growth did not vary between P14, M10, M10.Ca and ADJ (Figure 2B,C). However, the highest nutrient medium M14 had decreased photoautotrophic growth, particularly in the first few days after inoculation (Figure 2B,C). Because M14 was growth-inhibiting for photoautotrophic cultures, one or more elements may have induced nutrient toxicity physiology in -Glc cultures. In the lowest nutrient medium P14, growth declined by the second day of +Glc treatment, confirming P14 was insufficient for maximum +Glc growth (Figure 2C). Altogether, these results suggest that the macronutrient concentrations of M10 were optimal for both photoautotrophic and +Glc growth.

Doubling Ca in M10.Ca did not change growth or the internal Ca concentration compared to M10, and thus M10 was determined to be sufficient for *C. zofingiensis* medium (Figure 2C). Interestingly, P14 cells showed internal cellular hyperaccumulation matched with spent medium depletion of exogenous Mn (Figure S2), which did not occur in any other culture. While growth was not significantly limited in -Glc P14 growth, the unique hyperaccumulation further suggested that P14 levels altered nutrient import physiology, potentially as a signaling response to low nutrient concentrations in P14.

ICP-MS measurements of spent media indicated concentration adjustments were required to maintain replete nutrient levels through +Glc logarithmic growth phase. While ADJ and the M10 medium had similar growth, the spent medium of ADJ after +Glc treatment had more significant macronutrient losses. For example, ∼73% of P and ∼25% of S was depleted in ADJ spent medium after three days of +Glc, while M10 medium was reduced ∼25% P and ∼9% of S after three days +Glc (Figure 2D). These losses were not reflected by a unique hyperaccumulation of S or P per biomass in ADJ and indicate that the initial S and P increases from Bristol’s to make ADJ were still not sufficiently replete (Figure 2E). The macronutrient supplement of M10 was thus determined as more appropriate than ADJ for maintaining +Glc cultures in a replete state. Since the M10 medium is designed to support, in three-fold excess, the biomass of cultures achieved after 5 days of Glc treatment, we anticipated cells with M10 levels of nutrients should remain physiologically replete of all measured nutrients for ∼5 days of continuous +Glc consumption.

### Optimizing pH, nitrogen, potassium, and sodium for replicability

After using ICP-MS to determine replete nutrient concentrations to support +Glc growth in the design of the base medium M10, we further optimized other environmental variables for consistent growth experiments.

First, we evaluated ideal pH and buffers that would enhance growth and maintain environmental replicability of M10 medium. Throughout the optimization, we found that *C. zofingiensis* grew better in slightly alkaline cultures (pH > 7.5) and had progressively lower growth with pH ≤ 7 (Figure S3A,B). Therefore, ADJ medium and M10 had been set to pH of 8.0 with phosphate as buffer (Table S1). However, across several experimental replicates we were unable to detect significant differences between pH 7.5 and pH 8.0 in photoautotrophic growth curves. To reduce the increase in nutrient precipitation effect that would occur with excess alkalinity and to use a pH amenable to the pK_a_ of common laboratory buffers (e.g. HEPES), we lowered M10 medium pH to 7.5.

We then tested buffers suited for maintaining pH 7.5 to ensure they had no effect on algal physiology compared to a control medium that was only buffered by K_2_HPO_4_/KH_2_PO_4,_ from which a significant amount is assimilated (Figure 2D), reducing the buffer capacity of the media during growth. Growth curves of 20 mM HEPES and Tris buffers and the phosphate control showed no difference in volumetric biomass growth (Figure S4A). However, Tris-treated cultures had significantly lower cell number and division rates, resulting in larger individual cells, compared to both HEPES and control cultures (Figure S3E,D). All growth parameters were not significantly different between HEPES and control (Figure S3E,D). Measurements of the pH of spent media showed that phosphate buffer control did not prevent the tendency of photoautotrophic cultures to become more alkaline with time, but HEPES buffer could maintain pH at ∼7.5 for 6 days of photoautotrophic growth. With its strong pH control and no indication of impact on physiology, we chose HEPES buffer to improve M10 medium’s (now called M10 HEPES, Table S1) environmental consistency through logarithmic growth.

We next assessed nitrogen levels and sources manually, as we could not detect this compound by ICP-MS. Nitrogen is considered the most abundant biological element after carbon, oxygen, and hydrogen (Merchant et al., 2020). While algae would invest less energy in nitrate reduction if they were provided ammonium (NH_4_^+^), we found that *C. zofingiensis* did not grow in concentrations ≥22.5 mM NH_4_Cl (Figure S4A,B). *C. zofingiensis* could also grow in 22.5 and 45 mM NaNO_3_ with no apparent grow defects. To ensure NO_3_^-^ levels could be both high and proportionally greater than any other nutrient in the medium, we increased M10 medium levels from 22.5 mM to 45 mM NO_3_^-^ (now called M10 HEPES N Boost, Table S1) to keep nitrogen in excess for replete nutrient growth and avoid nitrogen deprivation in +Glc.

K and Na internal concentrations could not be matched to the optimized ratio of elements of cells due to their co-occurrence in the salts used to prepare the medium (e.g. NaNO_3_, K_n_H_n_PO_4_). Several plant studies show that the ratio of Na/K ions is important in salinity responses (Schachtman and Liu, 1999). The ratio of Na/K in this defined medium could be best adjusted in the phosphate and nitrate that is provided to the cells. To ensure cellular health, we compared a gradient of different Na/K ratios from 1:5 to 5:1 and found no significant impact on growth rates (Figure S5). Rather than requiring additional steps to incorporate various K/Na salts, our medium recipes only use single sources of NaNO_3_ and K_n_H_n_PO_4_ salts.

### Adjusting iron for a consistent intracellular ionome in logarithmic growth

Finally, we adjusted the medium to ensure the intracellular ionome was consistent through logarithmic growth phase, which is essential to reduce time-dependent confounding effects in time course studies. As hyperaccumulation can be a feature of nutrient excess (Hui et al., 2022), we evaluated the relationship between external concentration and the intracellular ionome levels of photoautotrophic cultures across the media variations and timepoints we obtained through ICP-MS data (Figure 3). Intracellular nutrient concentrations were normalized to moles of S, a method to standardize comparisons across different experiments (Schmollinger et al., 2021; Hui et al., 2022). Promisingly, most elements had consistent concentration per S despite ≥10-fold external range concentration of each element, showing external concentration usually does not impact internal nutrient import (Figure 3). However, iron was the largest outlier, showing increasing internal concentration and variability as its external concentration exceeded 100 µM. To a lesser extent, intracellular copper also increased with external concentration, but the mean increase plateaued above 10 µM external Cu, suggesting Cu quotas were saturated.

**Figure 3.**
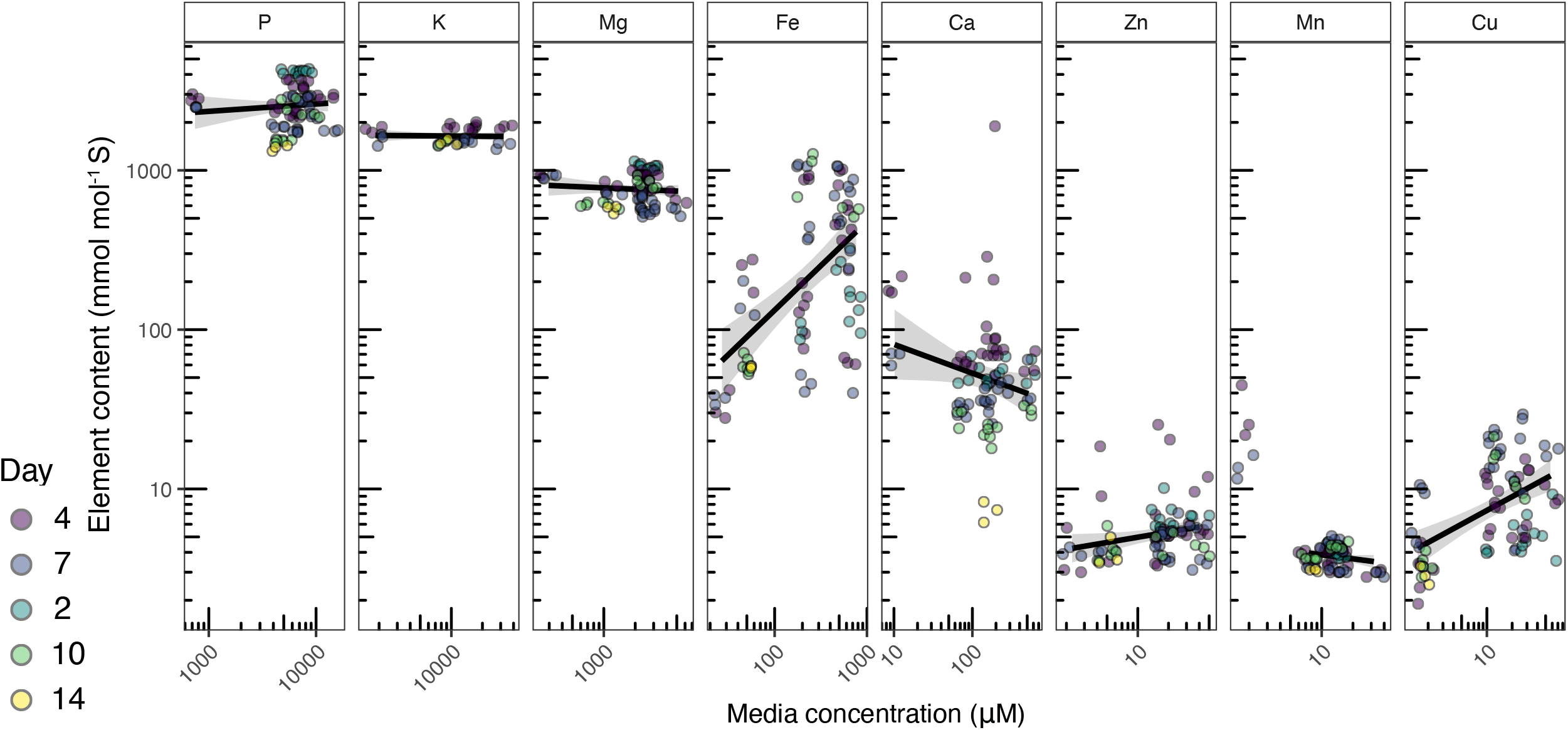
Intracellular nutrient levels across different external nutrient concentrations. Intracellular ionome variation per element normalized by cellular sulfur concentration across 81 photoautotrophic samples under diverse media regimes and sampling time points (point color represents day after inoculation). The fresh medium concentration of each element per samples is represented on the *x*-axis of each panel, whose range varies according to nutrient. The black line represents the linear regression conducted on internal elemental concentration as a function of medium concentration. For Mn, 12 P14 biological samples were plotted but not considered in the linear regression as they represented a hyperaccumulation in low external nutrient concentration that was not reflected in the range of higher concentration (Figure S2). Grey regions represent 95% confidence interval of regression model (*n*=81).

We compared iron quotas across *C. reinhardtii* and *C. zofingiensis* to determine that this nutrient responsive pattern was not due to *C. zofingiensis* simply requiring very high iron quotas. The M10 medium predicted a need for ∼210 µM Fe for biomass growth, and the determined the replete iron quota is 20 µM Fe for *C. reinhardtii* (Kropat et al., 2011). Furthermore, at similar concentrations to M10 medium (200 µM), *C. reinhardtii* hyperaccumulates iron in an excess state with no obvious impact on physiology or growth in standard conditions (Long and Merchant, 2008; Hui et al., 2022). Therefore, we suspected the media concentration-dependence of internal iron accumulation was due to a conserved green algal excess iron response and not due to specifically high iron needs in *C. zofingiensis*.

We tested if Fe concentration could be lowered to create a consistent ionome at logarithmic growth phase. Cells were grown in M10 medium with 50 or 200 µM Fe and all other nutrient concentrations remained constant. No significant growth change occurred between iron treatments through photoautotrophic growth or after +Glc treatment (Figure 4A,B). The intracellular ionome variation was sampled throughout photoautotrophic log phase (days 3-5). All elements including Fe had consistent logarithmic growth phase ionomes at 50 µM Fe (Figure 4C). However, Fe was the only element with log phase accumulation differences at 200 µM Fe (Figure 4), where it was higher and had greater variation at 200 µM Fe compared 50 µM Fe (Figure 4D). To create a consistent intracellular ionome, we adjusted the final medium to 50 µM Fe, avoiding both deficiency and excess nutrient impacts of *C. zofingiensis* during photoautotrophic and +Glc logarithmic growth. Based on using ionomics to define the optimized ratio of nutrients for replete growth, we named the final growth medium *<u>C</u>hromochloris zofingiensis* <u>O</u>ptimized <u>R</u>atio of <u>E</u>lements or CORE medium.

**Figure 4.**
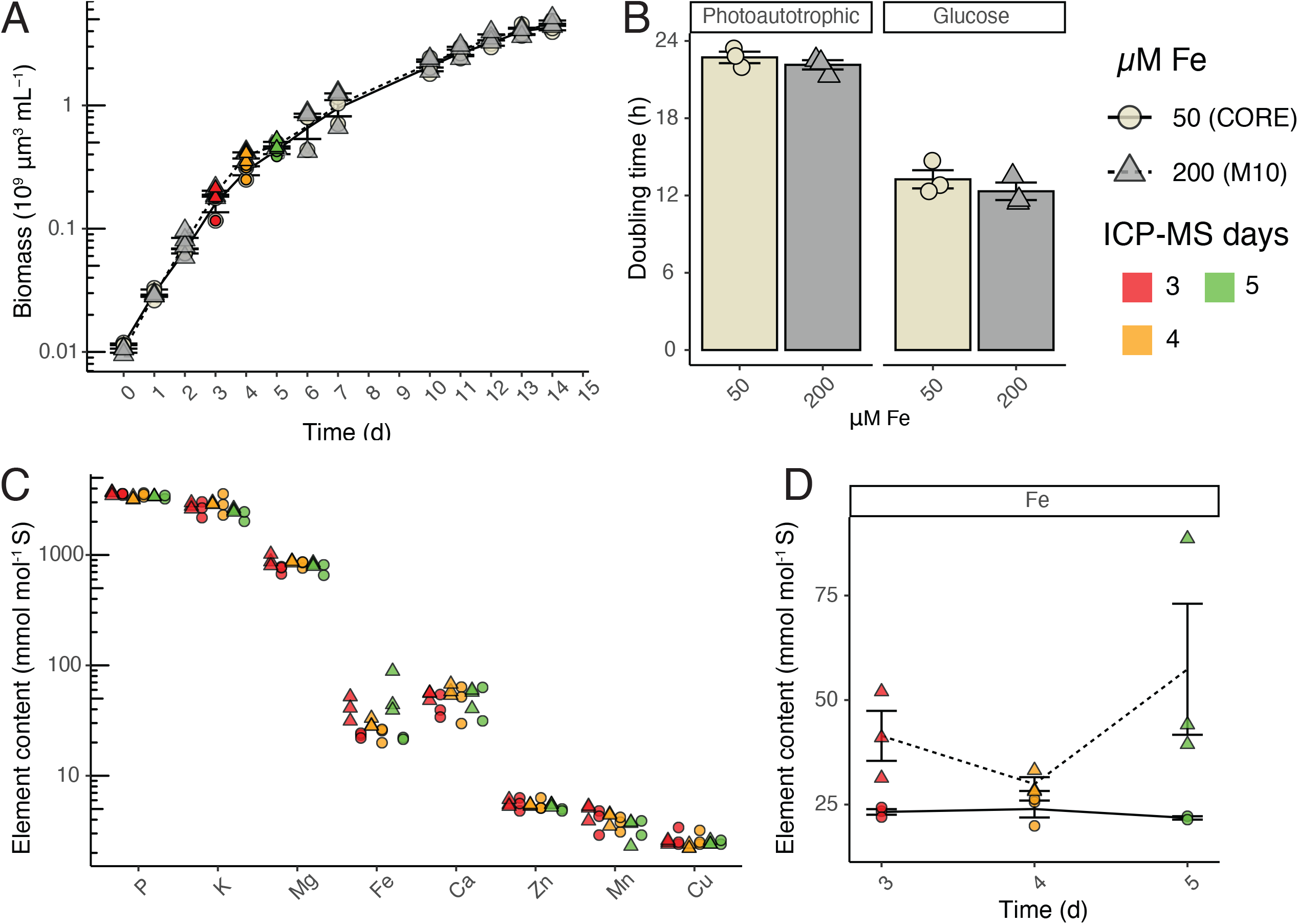
A consistent intracellular ionome is maintained through photoautotrophic logarithmic growth. **(A)** Photoautotrophic volumetric biomass growth curve of M10 HEPES N Boost medium (Table S1) with 200 µM Fe (grey, triangle, dotted line) vs. 50 µM Fe (beige, circles, solid line). Days 3, 4, and 5 samples were collected for intracellular ionomic quantification and are overlayed with red, orange, and green, respectively. **(B)** Volumetric biomass doubling time (h) of 50 vs. 200 µM Fe treatments in -Glc and +Glc (140 mM Glc, single dose). **(C)** The elemental content (normalized by cellular sulfur concentration) of all measured elements through log phase between 50 (circles) vs. 200 (triangles) µM Fe across days 3, 4, and 5. **(D)** Close-up of internal iron concentration across 50 vs. 200 µM Fe across days 3, 4, and 5. Because of this result, the 50 µM Fe medium represents the finalized CORE medium from this study. Line and bar graphs represent mean and error bars represent standard error (*n* = 2-3).

The final recipe for CORE medium can be found in Appendix S1 and can be used for controlled experiments comparing +Glc and -Glc physiology as well as a reference point to derive nutrient deficiency parameters and could be applicable to other microalgae particularly under mixotrophic or heterotrophic growth.

## Discussion

CORE medium was designed based on the internal ionome of *C. zofingiensis* to budget for +Glc growth while avoiding nutrient toxicity and maintaining replicability of log-phase cultures. Balancing the trade-off of limitation vs. toxicity makes CORE a suitable reference to study how nutrient deficiency acts in -Glc vs. +Glc cultures. Experiments aiming to understand long-term +Glc cultures, beyond the 5-day window supported by CORE, may require chemostats or turbidostats as more appropriate controlled systems. However, as it is unfeasible to have the many chemostats for experiments that require high sample number (such as combinatorial nutrient time courses (Jeffers et al., 2024), CORE medium is appropriate for multiomic investigations of nutritional status and their interplay. CORE medium may also be useful for other microalgae species that achieve high biomass, particularly through mixotrophic or heterotrophic growth. We have successfully grown another oleaginous alga, *Auxenochlorella protothecoides* (from class Trebouxiophyceae), in CORE medium with slight modifications based on species-specific physiology, despite being evolutionarily distant from *C. zofingiensis* (from class Chlorophyceae). Altogether, having a defined, nutrient-replete medium to support high biomass and bioproducts is essential for understanding these green algae.

Large macronutrient (N, P, K, Mg, S) supplements to the original Bristol’s medium were required to maintain replete concentrations in +Glc growth. Analysis of spent medium from +Glc cultures showed that S and P were highly consumed, indicating these elements are in specific demand in +Glc growth. Our previous research indicates Glc induces sulfur-related biosynthetic pathways (METE, methionine synthase; THIC and THI1/4 for thiazole biosynthesis) in *C. zofingiensis*, particularly during a switch-off of photosynthesis (Jeffers et al., 2024). +Glc also upregulates *de novo* fatty acid biosynthesis, which requires the S-containing coenzyme A (Roth et al., 2019a; Jeffers et al., 2024). These highly expressed pathways could drive increased S-uptake and demand in +Glc (Jeffers et al., 2024). Likewise, increases in the major P sink of carbohydrate phosphorylation and rRNA synthesis could underlie the higher P uptake in +Glc (Sulpice et al., 2014). As these S- and P-linked metabolic processes are connected to lipid and carbohydrate metabolism and biomass, their connection to +Glc signaling should be considered for metabolic engineering and design of large-scale bioprospecting of lipids.

Measuring the ionome of cells grown in nutrient compositions of a range of concentrations, showed that across the tested conditions, the relative internal compositions of most mineral nutrients is not influenced by increasing external concentrations. Fe is an exception, as its internal cellular concentration generally increased with external concentration. This pattern of *C. zofingiensis* excess iron accumulation is consistent with the *C. reinhardtii* iron accumulation (Long and Merchant, 2008; Schmollinger et al., 2021; Hui et al., 2022), where both organisms show accumulation patterns at ∼200 µM Fe. Like *C. zofingiensis*, iron hyperaccumulation in *C. reinhardtii* does not impact growth under standard conditions (Figure 4A,B; Long and Merchant, 2008; Hui et al., 2022). However, *C. reinhardtii* cells with hyperaccumulated iron are sensitive to high light treatment compared to replete Fe treated cells (Long and Merchant, 2008). Understanding accumulation patterns should be considered for media design across species. Beyond ensuring experimental replicability of the internal ionome (Hui et. al 2022), avoiding elemental accumulation in a medium will prevent confounding influences of non-nutrient related studies. For instance, researchers studying high light or reactive oxygen species response may want to ensure observed stress responses are not due to accumulated micronutrients exacerbating the impacts of the stress.

Measuring spent media and internal concentration of elements in algal biomass by ICP-MS, as pioneered by Kropat et al. (2011), creates a framework to discover elemental demands and identify an ideal standardized nutrient composition. From this reference medium, all other nutrient regimes and physiologies can be determined. The considerations of this technique may be optimized in industrial contexts to improve production costs. Biofuel production is often hindered by the costs of nutrients and the biomass loss that occurs during most green algal lipid induction by nutrient deprivation (Zhang et al., 2021; Ma et al., 2022). For industrial scale biofuel production, ionomic measurements of spent media and internal ionome may be calculated with its impact on biofuel and carotenoid production. This approach may be a key strategy to improve the yield and cost efficiency of valuable bioproducts. Therefore, consideration of algal optimal nutrient needs could serve as a reference point to find metabolic pathways (bioengineering targets) and nutrient adjustments (economic targets) that improve the sustainability of algae as viable replacements for fossil fuels.

While media rich with nutrients and carbon sources may produce high levels of biofuels and bioproducts, the nutrient and exogenous carbon supplements can be costly. Fortunately, microalgae can capitalize on nutrients and carbon sources from waste streams to reduce costs. The most promising source of carbon may be byproducts from table sugar manufacturing, which can be abundant with glucose, sucrose, and fructose (Liu et al., 2012). *C. zofingiensis* has been shown to accumulate biomass, lipids, and astaxanthin when grown with industrial waste from cane molasses (Liu et al., 2012). Furthermore, *C. zofingiensis* has been noted for its superior capabilities for both nitrogen and phosphorus uptake, showing that this alga can simultaneously treat wastewater while accumulating bioproducts (Zhao et al., 2018). Models have also shown competitive biodiesel prices using waste streams and a combination of microalgae (*C. reinhardtii*) and yeast (Gomez et al., 2016). With the understanding from this study of the nutrient profile that leads to optimal biomass, lipid, and astaxanthin accumulation, waste streams can be orchestrated for efficient and economically feasible bioproduction and wastewater treatment.

The CORE medium, designed based on the cellular ionome of mixotrophic cells, establishes a defined, nutrient-replete standard for future studies focused on *C. zofingiensis*. This medium is optimized for fast growth and can support high biomass and production of triacylglycerols in *C. zofingiensis* and can be used in mixotrophic and heterotrophic growth of other microalgae. Additionally, this standardized medium allows for controlled comparisons between experiments and laboratories and could be useful studies that investigate interplay of energy metabolism with nutrient demands and cofactors. This study also provides a strategy to improve yield and economic feasibility of microalgae and a nutrient profile to match from waste streams to improve cost efficiency of algal bioproducts.

## Experimental Procedures

### Media concentrations

Expanding from the initial medium described in Roth et al. (2019a), *C. zofingiensis* SAG 211-14 cultures were grown in iterative heterotrophic (+Glc) media recipes that were altered based on improvements to biomass, photosynthesis, and chlorophyll production from the Proteose medium (Table S1). ADJ medium, the base medium for this study, was composed of adjustments to Bristol’s macronutrient base (Bold, 1949). Stock solutions of metal micronutrients were complexed to Na_2_-EDTA as described by Kropat et al., 2011 (see Appendix S1 for details). Like in *C. reinhardtii* (Kropat et al., 2011), we did not detect internal concentrations of cobalt or boron in *C. zofingiensis* or find any impact of these elements on growth; therefore we excluded these elements starting with ADJ medium. In Figure 1, ADJ with 0.6 mM MgSO_4_ and 1.2 mM MgSO_4_ was tested simultaneously with no significant growth differences and was combined in student’s *t-* test of photoautotrophy vs. mixotrophy. Because sulfur had higher internal concentrations than its bound cation Mg, an additional K_2_SO_4_ supplement was added to the medium.

The finalized CORE medium contains 45 mM NaNO_3,_ 2.5 mM MgSO_4_, 1.5 mM K_2_SO_4_, 0.08 mM CaCl_2_, 5.7 mM K_2_HPO_4_, and 0.3 mM KH_2_PO_4_ (Table S1, Appendix S1). Micronutrients included 50 µM Fe-EDTA, 15 µM Mn-EDTA, 12 µM Cu-EDTA, 17.5 µM Zn-EDTA. Selenium and molybdenum sources were 0.03 µM Na_2_SeO_3_ and 0.057 µM (NH_4_)_6_Mo_7_O_24_. pH was altered to 7.5 and maintained with 20 mM HEPES. The final recipe is included in Appendix S1. In addition, all media concentrations used in the media optimization process are found in Table S1.

### Culturing and growth measurements

For all growth and nutrient optimizations, a starter culture of *C. zofingiensis* SAG 211-14 was inoculated at ∼1 × 10^5^ cells mL^-1^ from agar plates grown in constant light. All test growth curves were diluted from the starter liquid culture to 1 × 10^5^ cells mL^-1^ on inoculation day. Cells were grown in 50 mL of culture in 250 mL plastic beakers with petri dish lids that provided gas exchange inside an Infors HT Multitron Pro growth chamber with 100 µmol photons m^-2^ s^-1^ constant light, 150 rpm shaking, 25°C, and approximately 80% controlled humidity. The appropriate glucose concentration was added from a 2 M stock solution at 4 or 5 days after inoculation.

Cell density and volumetric biomass were assessed daily by a Multisizer 3 (Beckman Coulter). Volumetric biomass was calculated and extracted from Coulter Counter files by an R script described in Jeffers et al. (2024). In one case of extremely high density cultures (Figure 1B, day 24 of +Glc growth), % cellular volume of culture out of total liquid was approximated from the centrifuged wet cellular pellet volume in a Corning 50 mL Conical Falcon Tube.

### Glucose concentration measurements

To measure external glucose in the spent medium, a small amount of media supernatant (∼0.5 mL) was separated from cells by centrifugation. Samples were either assayed immediately according to the Megazyme D-Glucose Assay Kit (K-GLUHKR) or placed in a -20°C freezer. On wet ice, a master mix was prepared fresh containing 50 mM HEPES (pH 7.5), 5 mM MgCl_2,_ 1.33 mM NADP^+^, 4 mM ATP. A standard curve of glucose concentrations from 0 to 1 mM was established in technical triplicates, diluted to a final 200 µl volume with master mix in a 96-well plate. Technical triplicates of each spent medium sample were added to wells and diluted with master mix to be between 0 and 1 mM glucose. Absorbance at 340 nm was measured, and then 2 µL of Megazyme hexokinase + glucose-6*-*phosphate dehydrogenase (“Reagent 2”) that was stored at 4°C was added to each well, and the change of absorbance in each well was assayed over 20 minutes using a Tecan Infinite M1000 Pro (Tecan, USA) and used to quantify glucose concentration.

### Quantifying elemental concentrations by inductively coupled plasma mass-spectrometry

4 to 10 mL of culture were transferred to a 50 mL falcon tube, which was centrifuged for 3 min at 3220 *g* in an Eppendorf 5810R centrifuge. A fraction (<0.5 mL) of the supernatant was saved for spent medium analyses and frozen at -20°C. The pellets were resuspended in ∼10 mL of 1 mM Na_2_EDTA (pH 8.0) solution for three consecutive wash/centrifugation steps to remove residual cell-associated nutrients. The final wash steps took place in 15 mL falcon tubes and included a final wash with 10 mL of millipore water to remove remnants of EDTA before the analysis. The cell pellets were frozen at -20°C. All glassware used to hold EDTA solutions was acid washed in 50% HCl as previously described (Glaesener et al., 2013).

Prior to ICP-MS/MS analysis, cell pellets were digested with 70% ICP-MS grade nitric acid, first at room temperature overnight, then followed by additional incubation at 65°C until the cell pellet was completely dissolved. Elemental concentrations were quantified for both cell pellet and spent medium in an Agilent 8900 Triple Quadrapole ICP-MS/MS using methods and standardization as described (Schmollinger et al., 2021). Ionic measurements were either normalized to the volumetric biomass of the pellet or by moles of sulfur.

### Statistical analysis

All *t-*tests and linear models were conducted in R (v. 4.1.1). Plots were produced with ggplot2.

### Data Availability Statement

All relevant data for the manuscript and its supporting materials can be found in the Open Science Framework (link: https://osf.io/5bjcf/).

## Acknowledgements

We thank Christopher Baker for sharing the original glucose assay protocol, which we adapted for *C. zofingiensis*, Madeli Castruita for sharing the EDTA wash protocol for preparing samples for ICP-MS, and Michelle Meagher for providing measurements of iron concentration in Proteose medium. We thank Dimitrios Camacho for thoughtful comments on the manuscript. This material is based upon work supported by the U.S. Department of Energy, Office of Science, Office of Biological and Environmental Research, under Award Number DE-SC0018301. This article is subject to HHMI’s Open Access to Publications policy. HHMI lab heads have previously granted a nonexclusive CC BY 4.0 license to the public and a sublicensable license to HHMI in their research articles. Pursuant to those licenses, the author-accepted manuscript of this article can be made freely available under a CC BY 4.0 license immediately upon publication. K.K.N. is an investigator of the Howard Hughes Medical Institute.

## Short Legends for Supporting Information

### Supplemental Figures

**Figure S1**. Increasing nitrogen with glucose improves visible chlorophyll production.

**Figure S2**. Manganese and magnesium hyperaccumulated in low nutrient medium.

**Figure S3**. pH and buffer impacts on culture growth.

**Figure S4**. Effect of nitrogen species and concentration on *C. zofingiensis* growth.

**Figure S5**. Effect of potassium to sodium ratio on *C. zofingiensis* growth.

**Supplemental Table S1** All media compositions used in the study through optimization.

**Appendix SI** Recipe for finalized CORE Medium.

