## Supporting Information for "An algal nutrient-replete, optimized medium for fast growth and high triacylglycerol accumulation"

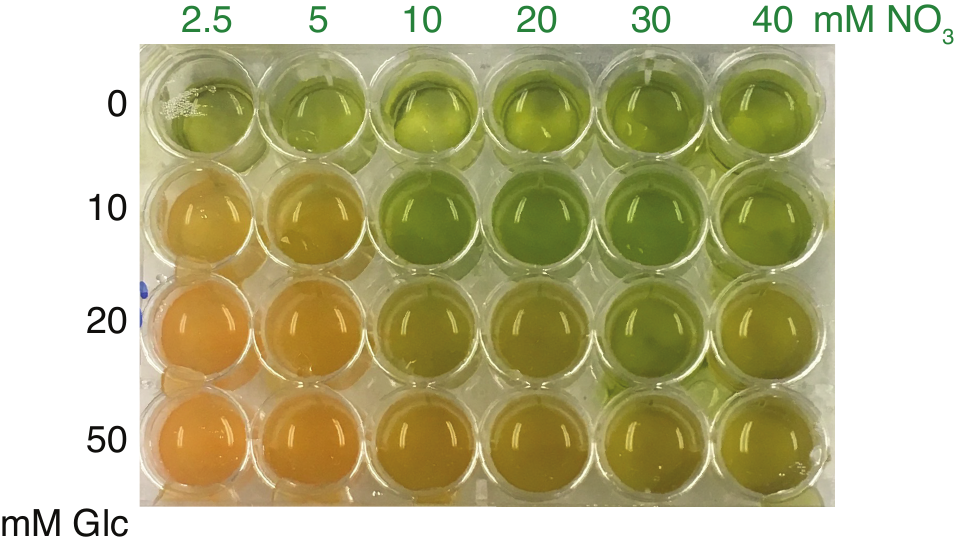

**Figure S1. Increasing nitrogen with glucose improves visible chlorophyll production.**

An example of cultures in a 24 well-plate resuspended across nitrate gradient (columns) and glucose gradient (rows). Photograph is taken 4 days after a seed culture was centrifuged and resuspended in the various N media with Glc treatment in each well. Cultures are grown in Bristols Medium (Bold 1949) with Hutner’s micronutrient supplement (Hutner et al., 1950) in 16 hour light:8 hour dark cycles at 100 µmol photons m^-2^ s^-1^. Glucose was only treated once in the cultures.

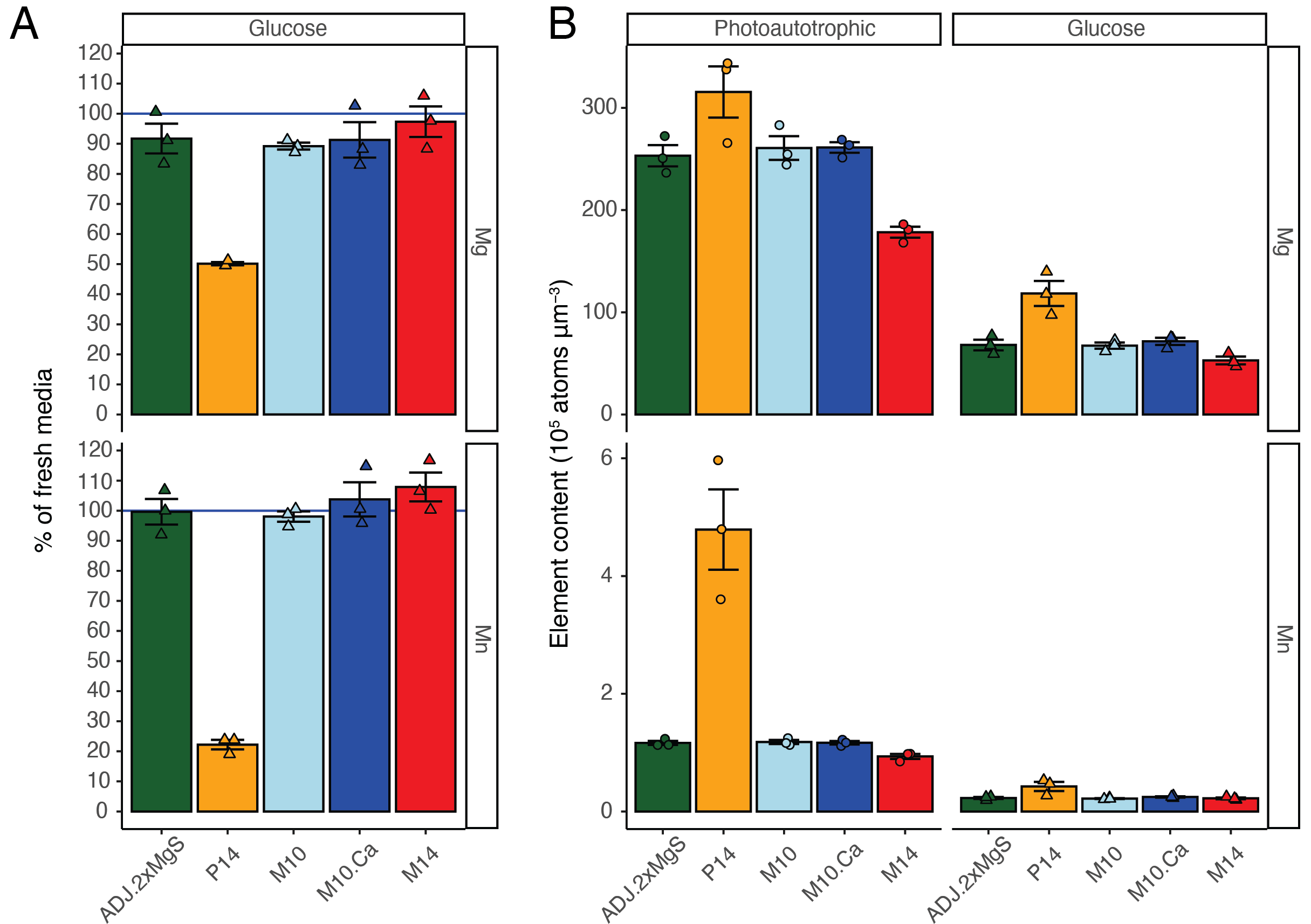

**Figure S2. Manganese and magnesium hyperaccumulated in low nutrient medium.**

**(A)** Percentage of magnesium (Mg) and manganese (Mn) left in spent medium on day 8 of +Glc growth of various media (*x-*axis). Depletion of Mn in P14 was already apparent by day 4. **(B)** Corresponding elemental hyperaccumulation of Mn in cells on day 7 for both photoautotrophic and +Glc samples in P14. Bar graphs represent mean and error bars represent standard error (*n* = 3).

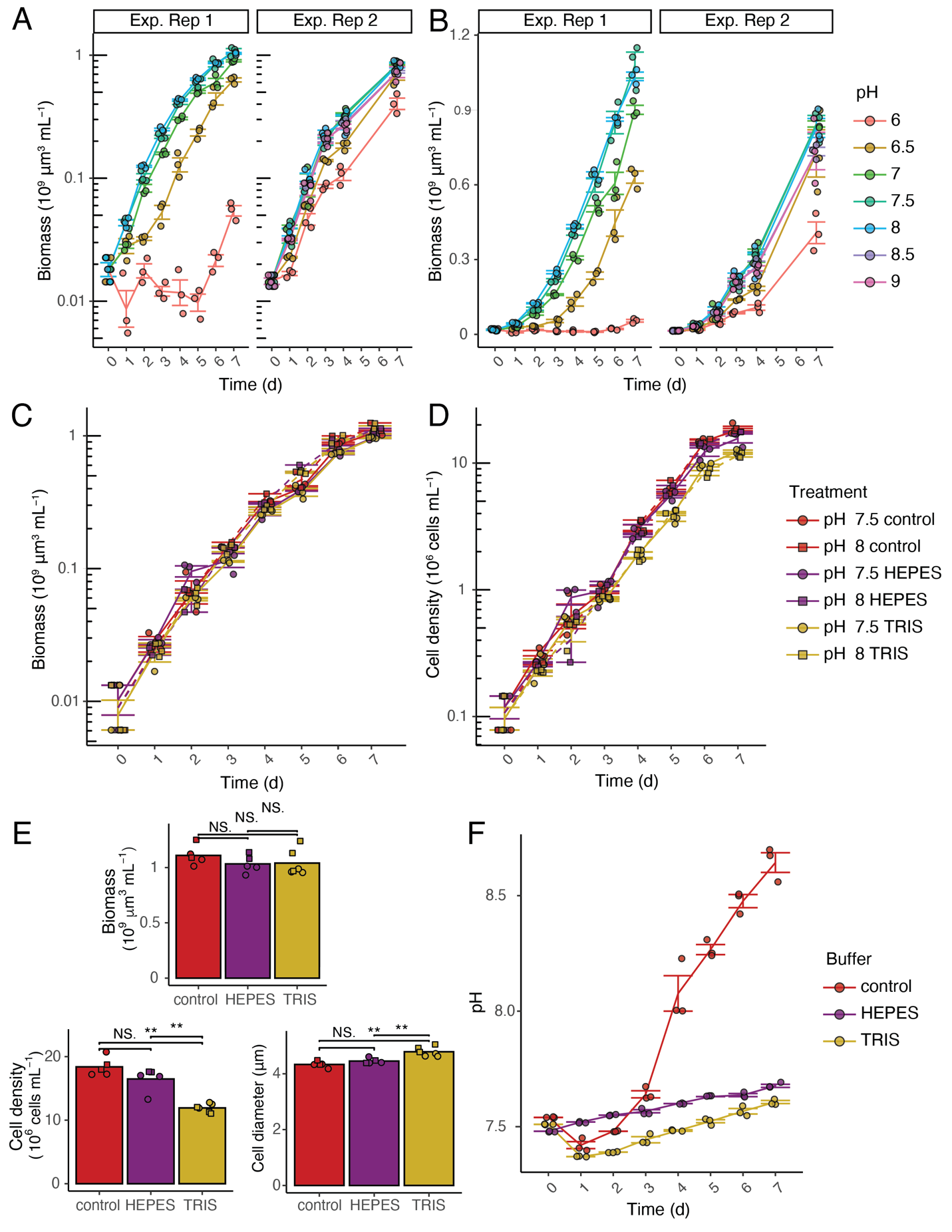

**Figure S3. pH and buffer impacts on culture growth.**

pH gradient volumetric biomass growth curves of two experimental replicates, conducted in M10 medium with only phosphorus buffer. For **(A)**, *y*-axis is in log scale while for the same data *y-*axis is not scaled in **(B)**. Colors refer to pH of media at the start of the experiment. Volumetric biomass **(C)** and cell density (**D)** growth curves of cultures with solely phosphate as a buffer (control, red), or with the addition of 20 mM HEPES (purple) or 20 mM Tris (gold). pH 7.5 (solid) and 8 (dashed) were also tested for each buffer treatment and were indistinguishable. **(E)** Day 7 measurements of volumetric biomass (top), cell density (lower left) and cell diameter (lower right). Statistics is done by pairwise *t-*test (N.S. refers to corrected *p-*value­ ­≥ 0.05 while ** refers to *p-*value < 0.01). **(F)** pH of media over time of three buffer treatments with media having a starting pH of 7.5. Line and bar graphs represent mean and error bars represent standard error (*n* = 3-6).

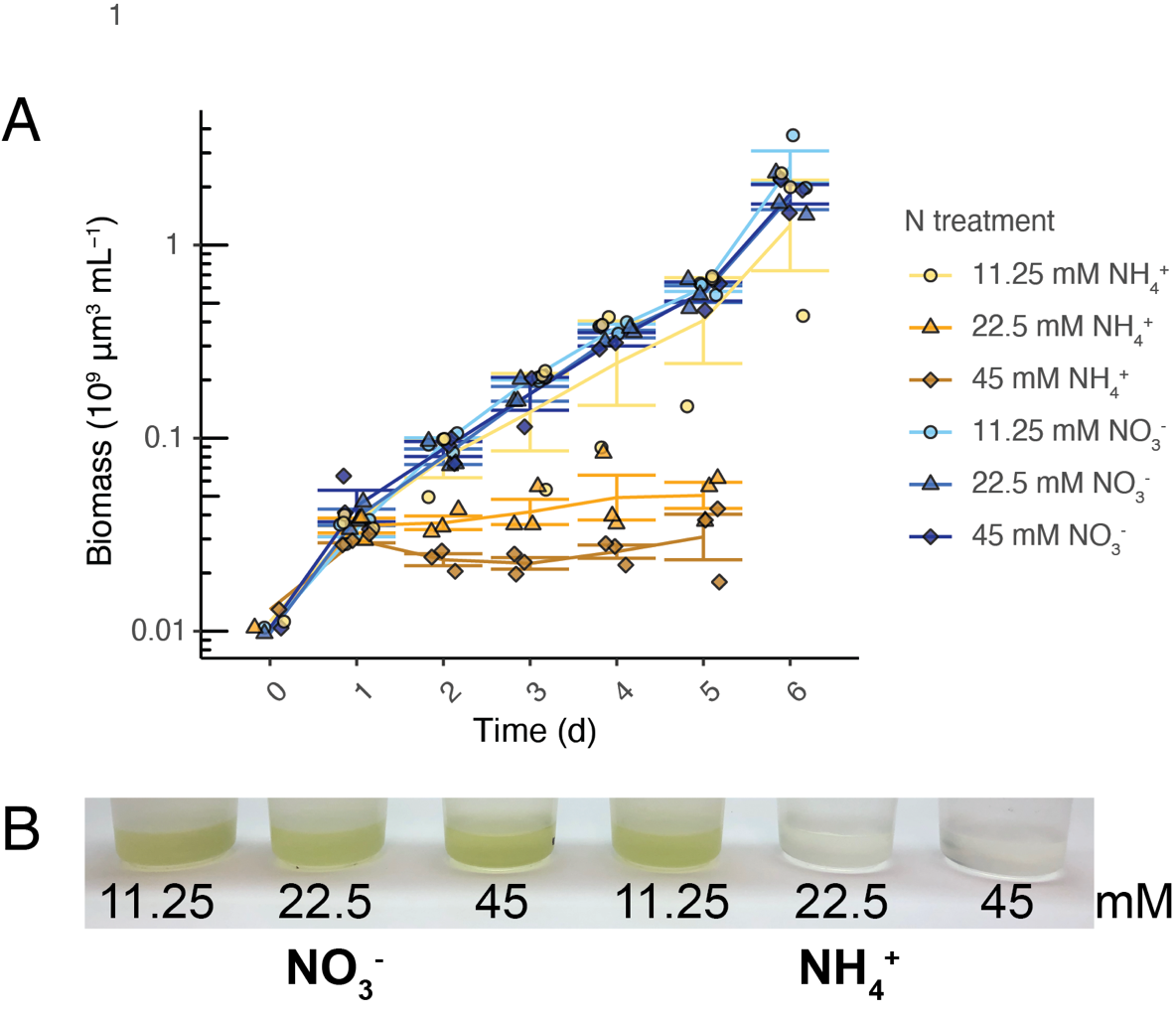

**Figure S4. Effect of nitrogen species and concentration on *C. zofingiensis* growth.**

**(A)** Volumetric biomass growth curves of cultures inoculated into different levels of ammonium and nitrate. Day 6 was not measured in ammonium cultures that failed to grow. **(B)** Photograph of autotrophic cultures of each treatment on day 3. Besides nitrogen, all other medium parameters pertain to M10 + 20mM HEPES solution (Table S1). Line graphs represent mean and error bars represent standard error (*n* = 2-3).

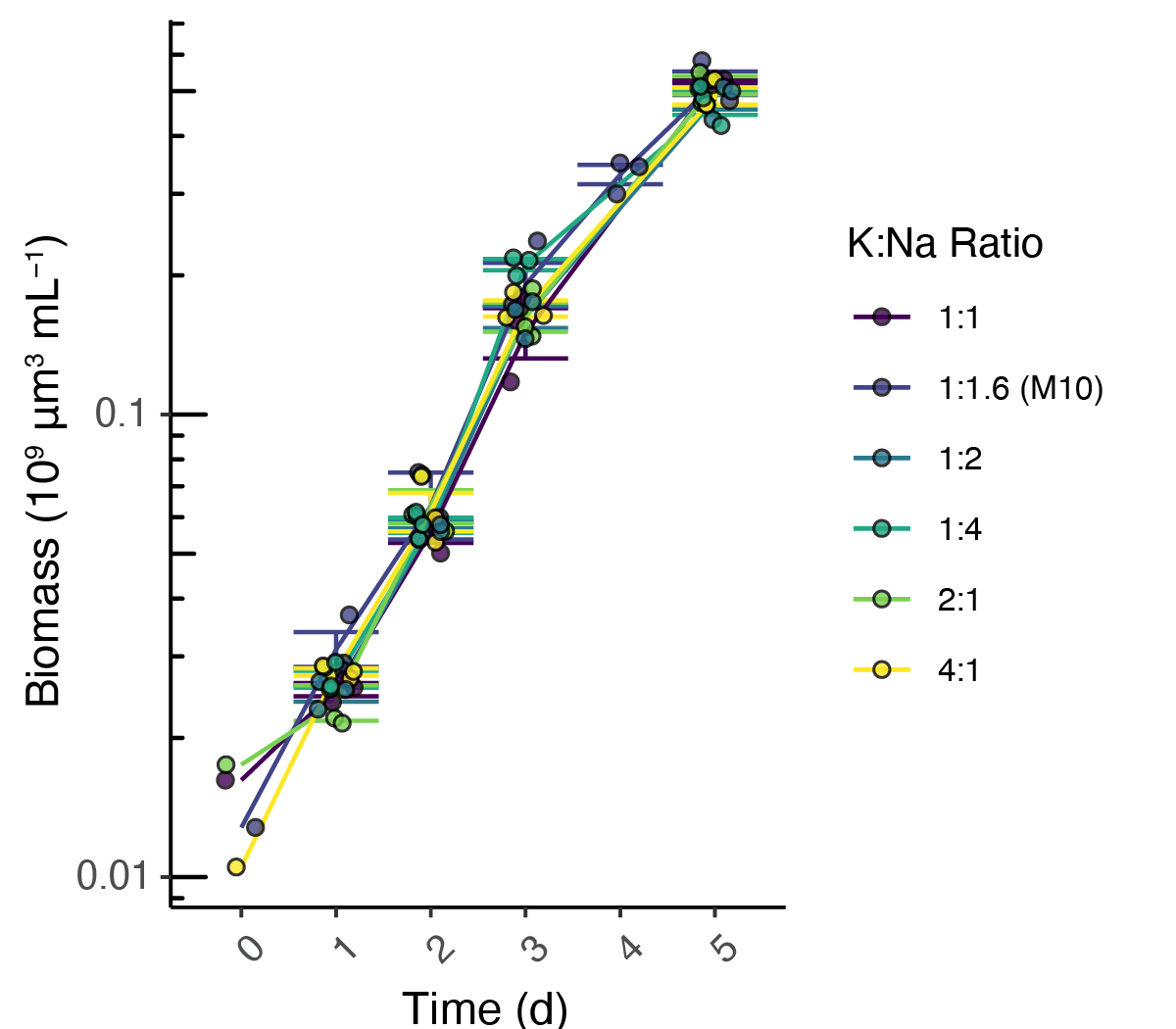

**Figure S5. Effect of potassium to sodium ratio on *C. zofingiensis* growth.**

Volumetric biomass growth curves of cultures where potassium to sodium concentration ratios (K/Na) are intentionally altered while the anions NO_3_^-1^ and PO_4_^-3^ are maintained at the same concentrations. “1:1.6 Ratio (CORE)” treatment indicates the CORE medium K/Na, which is the result of using only sodium nitrate and potassium-phosphate as salt sources in the medium. Beside K and Na, all other medium parameters pertain to M10 medium (Table S1, Appendix I). Line graphs represent mean and error bars represent standard error (*n* = 2-3).
