## Appendix S1 for "An algal nutrient-replete, optimized medium for fast growth and high triacylglycerol accumulation"

**Recipe:**

**CORE (*Chromochloris*-Optimized Ratio of Elements) Medium**

***with* HEPES buffer**

***Recommendations***

***Potassium phosphate solutions need to be autoclaved in a separate bottle from the other nutrients and recombined once they are at room temperature—the solution will precipitate otherwise. When making plates, autoclave the agar with the non-phosphate media; with the phosphates, it looks caramelized and studies show this combination can produce H_2_O_2_ that inhibits microbial growth*

***HEPES is recommended to be filter sterilized and added after autoclaving. We use a 1M HEPES solution at pH 7.5, and just add 20 mL to 480 mL of Phosphate stock*

**1 L of CORE Medium**

1. *In 1 L graduated cylinder add 10 mL of each of the “-P Macronutrients”*

| **Stock** | **Stock Solutions (mM)** | **Final Conc. (mM)** | **mL of stock** |
| --- | --- | --- | --- |
| **NaNO3** | 2250 | 45 mM | **20.00** |
| **MgSO4** | 250 | 2.50 mM | **10.00** |
| **CaCl2** | 8 | 0.08 mM | **10.00** |
| **K2SO4** | 150 | 1.5 mM | **10.00** |

*also add corresponding volumes of micronutrients :*

*Micronutrients + 1 mL of 25 mM EDTA*

| **Stock** | **Final Conc.** | **(mL) of stock** |
| --- | --- | --- |
| **Fe-EDTA** | 50 µM | **2.50** |
| **Mn-EDTA** | 15 µM | **2.50** |
| **Cu-EDTA** | 12 µM | **6.00** |
| **Zn-EDTA** | 17.5 µM | **7.00** |
| **Se** | 0.03 µM | **0.25** |
| **Mo** | 0.057 µM | **2.00** |
| **Stock** |  | **mL** |
| EDTA-Na_2_ | 25 µM | **1** |

**2.** Fill to 500 mL with Millipore H_2_O, and transfer to 1 L bottle.

3. ***optional*** Add 15 grams of agar to this salt stock (not the phosphate bottole).

4. In another graduated cylinder, add the 10 mL of the two potassium phosphates sources. Note a much higher concentration of dibasic to make the final pH closer to 7.5.

| **Stock** | **Stock Solutions (mM)** | **Final Conc. (mM)** | **mL of stock** |
| --- | --- | --- | --- |
| **K2HPO4** | 570 | 5.70 | **10.00** |
| **KH2PO4** | 30 | 0.30 | **10.00** |
| [Tot P] |  | 6.00 |  |

5. Fill to 480 mL* with Millipore H_2_O, put in a separate 1L bottle. ***You want to save room for sterile 20 mL HEPES buffer to be added after autoclaving***

**6. To autoclave:** Put both bottles in a small autoclave bin and fill with one inch of water*.* Autoclave for 40 min.

7. Once solution has cooled to room temperature, add the KP autoclaved solution to the other nutrients solution. Add 20 mM of HEPES 1 M, pH 7.5 to the solution. Label the media “CORE medium”). Pour into plates if making solid medium.

**Stock solutions:**

**Macronutrients (modified extensively from Bristols Media)**

****** acid wash these vessels to be amenable to metal deficiency studies**

(HCl-based acid-washing protocol can be found in Glaesener et al., 2013)

| **v** | **Stock Solutions (mM)** | **Stock #** | **g/L for Stock** | **Molecular Weight (g/mol)** |
| --- | --- | --- | --- | --- |
| **NaNO_3_** | 2250 | Sigma S343 | **191.23** | 84.99 |
| **MgSO_4 *_ 7 H_2_O** | 250 | Fisher S24514 | **61.62** | 246.48 |
| **CaCl_2_ * 2 H_2_O** | 8 | CD0050 | **1.18** | 147.02 |
| **K_2_HPO4** | 570 | PX 1570-1 | **99.28** | 174.18 |
| **KH_2_PO4** | 30 | Fisher BP362 | **4.08** | 136.08 |
| **K2SO4** | 150 | Sigma P9458 | **26.14** | 174.25 |

**Micronutrients:**

Micronutrient Recipes are from Kropat et al., 2011

****Acid wash the vessels that will contain these media before preparing ***
(HCl-based acid-washing protocol can be found in Glaesener et al., 2013)

**A. Preliminary concentrated stock solutions**

Pre-1. EDTA-Na_2_ concentrate	125 mM	13.959 g in ~ 250 ml, titrate to pH 8.0 with

trace element grade KOH (~1.7 g), and bring up to a volume of 300 ml

Pre-2. (NH_4_)_6_Mo_7_O_24_	concentrate	285 µM	(NH_4_)_6_Mo_7_O_24_⋅4H_2_O: 0.088 g, bring up to a

volume of 250 mL

Pre-3. Na_2_SeO_3_ concentrate	1 mM	Na_2_SeO_3_: 0.043 g, bring up to a volume of

250 mL

**B. Individual Stock Solutions for medium (1000×)**

| Stock Solution | Concentration  in stock | Composition |
| --- | --- | --- |

1. EDTA-Na_2_	25 mM	EDTA-Na_2_: 50 mL of 125 mM EDTA-Na_2_ concentrate

(Pre-1) from Step A

2. (NH_4_)_6_Mo_7_O_24_	28.5 µM*	(NH_4_)_6_Mo_7_O_24_⋅4H_2_O: 25 mL of 285 µM (NH_4_)_6_Mo_7_O_24_

concentrate (Pre-2) from Step A

3. Na_2_SeO_3_	0.1 mM	Na_2_SeO_3_: 25 mL of 1 mM Na_2_SeO_3_ concentrate (Pre-3)

from Step A

4. Zn⋅EDTA	2.5 mM	ZnSO_4_⋅7H_2_O: 0.18 g

2.75 mM	EDTA-Na_2_: 5.5 mL of 125 mM EDTA-Na_2_ concentrate

(Pre-1) from Step A

5. Mn⋅EDTA	6 mM	MnCl_2_⋅4H_2_O: 0.297 g

6 mM	EDTA-Na_2_: 12 mL of 125 mM EDTA-Na_2_ concentrate

(Pre-1) from Step A

6. Fe⋅EDTA	20 mM	FeCl_3_⋅6H_2_O: 1.35 g

22 mM	EDTA-Na_2_: 2.05 g

22 mM	Na_2_CO_3_ (sodium carbonate): 0.58 g

(Combine EDTA-Na_2_ with sodium carbonate in water and mix. Add

FeCl_3_⋅6H_2_O after the first two components dissolve. Do Not Use Pre-1.)

7. Cu⋅EDTA	2 mM	CuCl_2_⋅2H_2_O: 0.085 g

2 mM	EDTA-Na_2_: 4 mL of 125 mM EDTA-Na_2_ concentrate
